# BAI-Net: Individualized Anatomical Cerebral Cartography using Graph Neural Network

**DOI:** 10.1101/2021.07.15.452577

**Authors:** Liang Ma, Yu Zhang, Hantian Zhang, Luqi Cheng, Junjie Zhuo, Weiyang Shi, Yuheng Lu, Wen Li, Zhengyi Yang, Jiaojian Wang, Lingzhong Fan, Tianzi Jiang

## Abstract

Brain atlas is an important tool in the diagnosis and treatment of neurological disorders. However, due to large variations in the organizational principles of individual brains, many challenges remain in clinical applications. Brain atlas individualization network (BAI-Net) is an algorithm that subdivides individual cerebral cortex into segregated areas using brain morphology and connectomes. BAI-Net integrates topological priors derived from a group atlas, adjusts the areal probability using the connectivity context derived from diffusion tractography, and provides reliable and explainable individualized brain parcels across multiple sessions and scanners. We demonstrate that BAI-Net outperforms the conventional iterative clustering approach by capturing significantly heritable topographic variations in individualized cartographies. The topographic variability of BAI-Net cartographies shows strong associations with individual variability in brain morphology, connectivity fingerprints and cognitive behaviors. This study provides a new framework for individualized brain cartography and paves the way of atlas-based precision medicine in clinical practice.

## Introduction

Brain atlas has been an important tool to understand the neural basis of human cognition. Neuroanatomists have built a variety of macro- and microanatomical atlases to depict cyto-, myelo- and receptor architectures using a few postmortem human brains ^1-7^. Recent advances in noninvasive neuroimaging techniques, such as magnetic resonance imaging (MRI), provide an opportunity to explore the anatomical and functional organization of the living human brain and to make subsequent cartographic explorations of the human cerebral cortex in a large population ^8-16^. However, the majority of current brain atlases focus on a group representative mapping of the cerebral cortex, but ignore the variations of individual brains in terms of areal size, location, spatial arrangement and connectivity patterns due to genetic and environmental influences ^15, 17^. The precise mapping of individual-specific topographic organization is a critical step towards better understanding the structural-functional relationship of the human brain underlies cognition and behavior ^18-20^ as well as for personalized localization diagnosis and treatment of neurological disorders ^21, 22^.

Traditional individualized cartography of cerebral cortex has relied on the linear and non-linear registration based on the structural images in the volume space or cortical surfaces ^23^. Modern machine learning algorithms provide analytic tools to align cortical areas using multimodal neuroimaging data, including structural and functional localizers ^24, 25^, as well as anatomical^26^ and functional connectomes^18-20^. As one of the most commonly used approaches to reconstruct human connectomes, diffusion tractography has offered exclusive tools to map anatomical connections in the human brain non-invasively^27^. However, the anatomical accuracy and biological meaning of diffusion tractography is still controversy nowadays ^28, 29, 30^, which may bias the areal delineation on individual brains when directly applying the diffusion tractography results in individualized cortical cartography ^26^.

To tackle these issues, we first employed a fiber-tract embedding approach that projects the whole-brain tractography maps to individual fiber-tract space by using TractSeg ^31^. The resulting connectivity fingerprint indicates the probability of the chosen major fiber tracts in the individual tractography map. The connectivity fingerprint approach has been widely used in neuroscience research and demonstrated high consistency not only across subjects but also between homologous areas cross species^32^, providing a substantial neural basis to reveal individual variations in anatomical connectomes. Besides, we applied two additional structural constraints on the individualized cartography model in order to precisely characterize the connectivity features from individual anatomical connectomes. The first structural constraint was the areal location priors derived from the group atlas, which provide a blueprint of the general organizational principles on individual brains and guide the individual-specific cartography by using generalized knowledge inferred from a large population rather than limited measures of a single subject ^13, 18-20, 25, 33-36^. Using such populational priors, we achieved robust delineation of cortical areas on individual brains under various scanning conditions, and at the same time improved the inter-subject alignment of individual topography which has been a common issue in the individualized cartography model ^20, 37^. As an important characteristic of human connectomes, the local continuity constraint suggests that adjacent cortical areas generally follow a similar neural pathway and connect to adjacent neurons in the target area. In order to implement such continuity constraint on individual anatomical connectomes, we employed the convolutional operations on the vertex-level graph constructed from individual cortical surfaces and trained various graph convolutional kernels to integrate the context information of connectivity fingerprints at different spatial ranges.

With these in mind, we developed a Brain Atlas Individualization Network (BAI-Net) for individual-specific cartography constrained by both populational priors of areal locations (e.g. Human Brainnetome Atlas ^14^) and the local continuity of individual connectomes. The BAI-Net method consists of three key steps, i.e. construction of individual brain graph, embedding of individual connectivity fingerprints and areal delineation using the local context of connectivity fingerprints. Specifically, a large-scale vertex-level graph (32k vertices per hemisphere) was first constructed from individual cortical surfaces. After that, a graph neural network (GNN) architecture was implemented to merge the topographical organization of individual brains with the connectivity context of anatomical fingerprints. Using a gradient descent optimization algorithm and trained over a large population, the resulting GNN representations learned the tradeoff between individual-specific topography and globally aligned organizational principles. We first investigated the reproducibility and robustness of BAI-Net individualized cartographies under various acquisition conditions including multiple sessions and different scanners. We further evaluated the interpretability of the topographic variability revealed by BAI-Net cartographies, e.g. association with individual variability in brain morphology, connectivity fingerprints and cognitive behaviors as well as its heritability in the twin population.

## Results

### BAI-Net individual-specific cartography of cerebral cortex

The BAI-Net method (***Fig. 1***) was evaluated using 969 subjects from the HCP dataset (including 100 unrelated subjects used for model training) and 74 repeated scans (consisting of 14 subjects) from MASiVar dataset. The detailed information about the datasets used in the evaluation steps was listed in ***Supplementary Figure S1***. During model training, a vertex-level brain graph was first constructed from T1-weighted images of individual brains, with each node indicating a vertex in the cortical surface and the edge indicating whether two vertices shared in a triangle in the cortical mesh. Then, the anatomical fingerprints, derived from the probabilistic tractography on the individual diffusion MRIs, were embedded as node/vertex features in the brain graph. Next, a graph neural network (GNN) was used to integrate the local context of the connectivity fingerprints and to update with a new representation that combines the topographic patterns in brain morphology and the context of connectivity fingerprints of the individual brains. Finally, the areal probability was inferred from the last layer of trained GNN.

**Figure 1.**
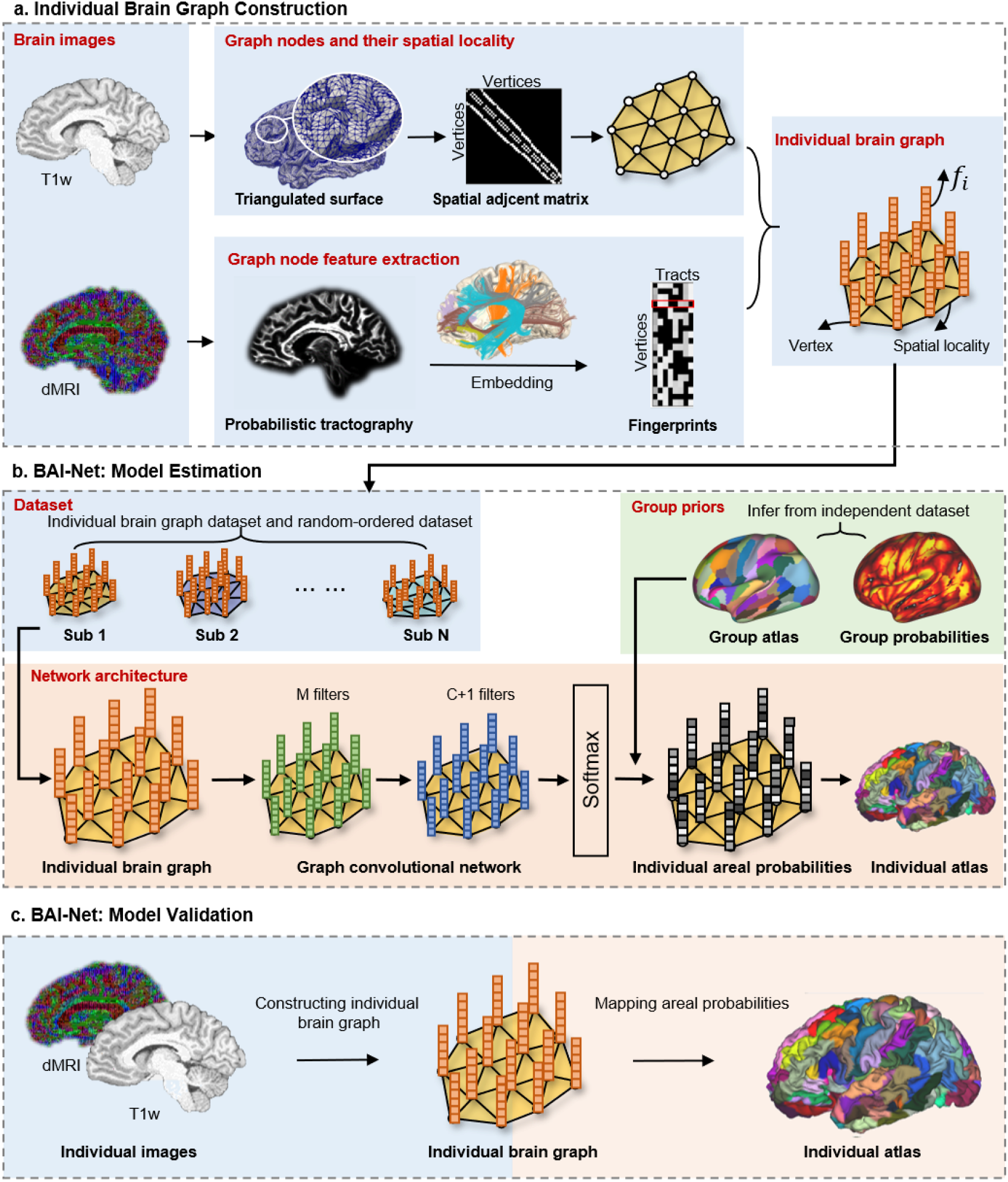
Schematic diagram of the Brain Atlas Individualization Network (BAI-Net) with group priors. a: Construction of individual brain graph. The cortical surfaces were reconstructed from the T1-weighted image using the Freesurfer and Connectome Workbench toolboxes. The individual brain graph was built based on the surface vertices, local edges, and connectivity fingerprints. b: BAI-Net: Model Estimation. Samples from the HCP training dataset and random-ordered dataset were fed as the inputs into the graph neural network. The outputs of the graph neural network for each node were areal probabilities. The group labels as well as the corresponding maximum probability maps of the group atlas were both registered from the group fs_LR32k surface to the individual surface tessellation. The network was optimized with probability-weighted loss function. c: BAI-Net: Model Validation. The regular step for cerebral cartography of a new subject was to build the individual connection graph (preprocessing, tracking, embedding, and normalization) and then to map it through trained network to get the areal probabilities as well as the individual cortical area with max probabilities.

### Largely retained global topographic pattern and considerable individual differences

The BAI-Net individualized cartographies generally followed a similar topographic pattern with the group atlas (average Dice score = 0.762±0.025 on the HCP test sets). The detailed maps of the areal borders and their overlaps with the group atlas were shown in ***Supplemental Fig. S2***. On one hand, the maximum probability map (MPM) and areal size of individualized cartographies were highly consistent with the group atlas (Dice=0.88). On the other hand, considerable individual differences were detected in the areas associated with high-order cognitive functions, for instance, the inferior frontal gyrus (IFG), inferior parietal lobe (IPL), middle temporal gyrus (MTL), and anterior cingulate cortex (ACC). Our results indicated that the BAI-Net individualized cartographies mostly retained the global topographic organization of cerebral cortex inherited from the group atlas, and uncovered considerable variations in the topographic arrangement of the association areas by aggregating the context of connectional architecture on individual brains.

### Reproducible individual-specific topography

The reproducibility of individual-specific topography was evaluated using the HCP test-retest datasets, consisting of 44 healthy subjects, who have collected structural, diffusion, and functional MRI data in two independent sessions. The topography similarities between intra-(between the test and retest sessions of the same subject) and inter-subject (between different subjects from the same session) pairs of individualized cartographies was evaluated by Dice score. As shown in ***Fig. 2a***, the BAI-Net cartography generated highly reproducible individual topography at the subject level (Dice = 0.901±0.014 for the whole cerebral cortex), while maintained high variability across subjects (Dice = 0.745±0.025), with significantly lower topographic similarity between subjects than within subjects (*p* values < 10^−50^). Examples of individualized cartography in the right hemisphere were shown in ***Fig. 2b*** (maps of the left hemisphere shown in ***Supplementary Fig. S3***). Compared to the canonical iterative clustering (IC) approach, BAI-Net cartography revealed higher gaps between intra- and inter-subject topographic similarity (0.15 and 0.09 respectively for BAI-Net and IC, see ***Supplementary Fig. S4*** for detailed information of IC) and consequently uncovered more subject-specific characteristics in brain topography. For instance, the identity of BAI-Net individualized cartography was successfully predicted in the HCP test-retest sessions (accuracy=100% when classifying subjects based on the maximum similarity in topography).

**Figure 2.**
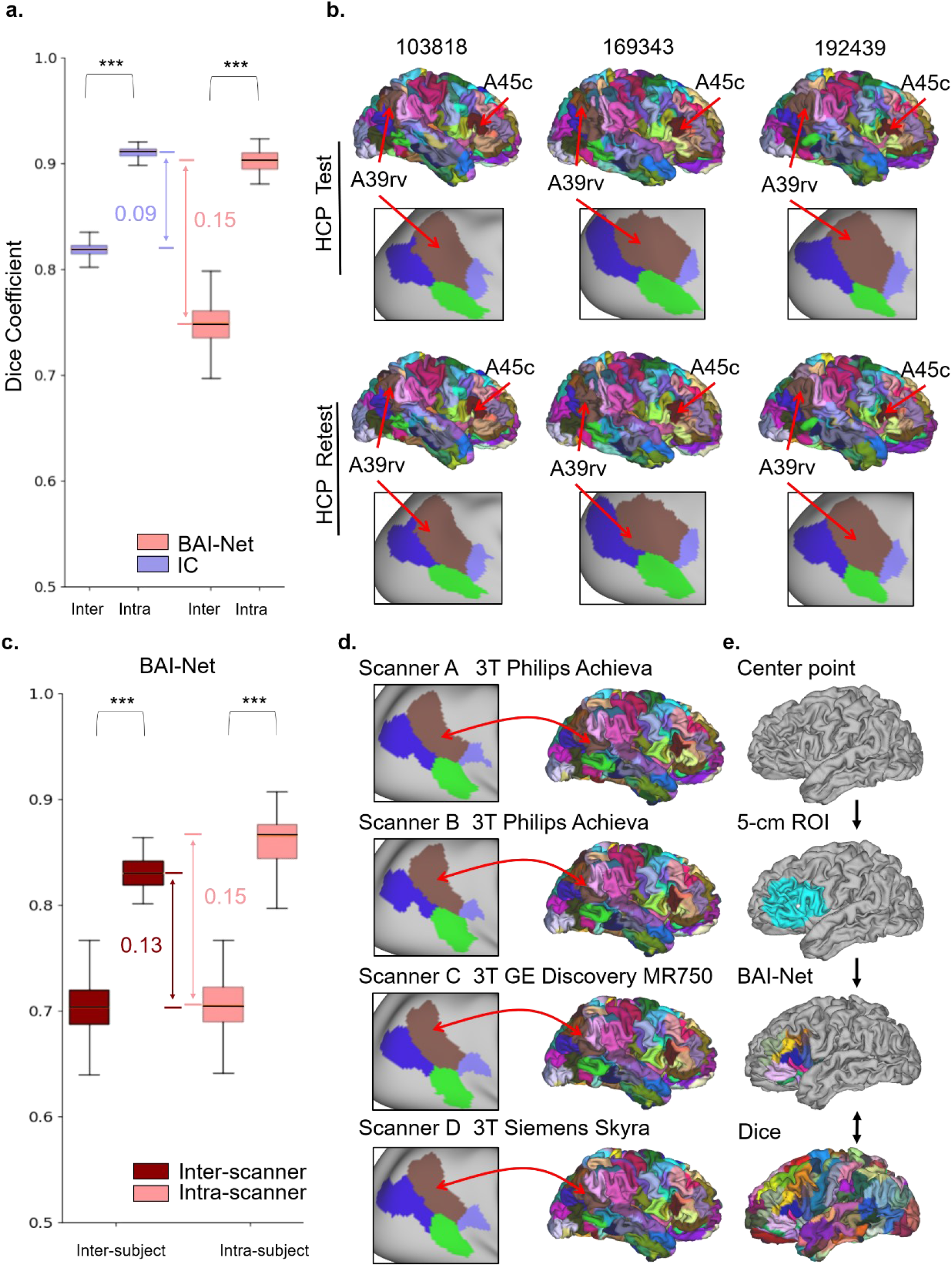
Reproducibility and specificity of BAI-Net individualized cartographies. a: Inter- and intra-subject variability of brain cartographys using BAI-Net and iterative clustering (IC) methods on the HCP test-retest dataset. b: Examples of BAI-Net individualized cartographies for three random subjects. c: Generalizability of the BAI-Net method on multiple scanners evaluated on the MASiVar dataset. d: BAI-Net individualized cartography for the same subject on four different scanners. e: Exemplar regional cartography when applying the BAI-Net method on a small region of interest, e.g. vlPFC. High overlaps were achieved between the regional cartography and whole-cortex cartography. The box represents the first and third quartiles in the distribution of the Dice scores.

### Robust performance across multiple scanners

The generalizability of the BAI-Net method was evaluated by applying the pre-trained HCP model onto the MASiVar dataset acquired from different scanners and sites. As shown in ***Fig.2c***, the BAI-Net cartography yielded much higher variability between subjects than within the same subject, not only within the same scanner (Dice score=0.860±0.024 and 0.707±0.027, respectively for intra-subject and inter-subject pairs), but also across different scanners (Dice score=0.830±0.015 and 0.703±0.025, respectively for intra-subject and inter-subject pairs). The BAI-Net cartography generated highly consistent cartographies of the same subject across four different scanners (***Fig. 2d***). It is worth noting that the reproducibility of individualized cartography was slightly lower in the multi-scanner dataset (MASiVar) than in HCP at both the intra- and inter-subject levels, mainly due to different scanning conditions in the two datasets. We further evaluated the inter-subject topography variability (ITV) by using Cohen’s *d* effect size. Compared to the IC method, BAI-Net method revealed relatively stable ITV for intra-scanner (Cohen’s *d* = 4.23) and inter-scanner cases (Cohen’s *d* =4.35) while the IC approach was more sensitive to the specific scanner (Cohen’s *d* = 6.64 and 3.31 respectively for within- and between-scanner ITV).

### Flexible and time-saving regional cartography

The BAI-Net method can be easily adjusted to the cartography of a small region of interest instead, namely the regional cartography. As shown in ***Fig. 2e***, the subdivision of the ventral lateral prefrontal cortex (vlPFC) was highly consistent with the whole-cortex cartography (Dice=0.92 for an exemplar subject, more examples can be found in ***Supplementary Fig. S5***). Another advantage of the BAI-Net regional cartography is the time-saving mode when applying to a small region. For instance, the BAI-Net cartography of vlPFC only takes 4 minutes, about one tenth of the time spending on the whole-cortex model (2.7k vs 32k vertices, respectively in the seed mask). In contrast, the IC method requires iteratively updating signals of the entire cerebral cortex at each iteration, which highly limits its applications on small regions and potentially biases the areal delineation when only local information was available (see ***Supplementary Fig S5b*** for three examples).

### Interpretability of BAI-Net cartographies

#### Topography variability associated with individual variability in brain connectomes and morphology

The effect size of the inter-subject topography variability (ITV) was evaluated by Cohen’s *d* on the HCP test-retest dataset, which computes the differences in individualized brain cartography between subjects after taking in account the intra-subject variability. The BAI-Net individualized cartographies exhibited large effect of ITV at the whole-brain level (Cohen’ *d* = 7.07 and 7.19 for the left and right hemispheres, respectively). The pattern of topography variability generally followed the functional and connectional gradient of the cortical organization (as shown in ***Fig. 3a***). Specifically, we found small ITV values in the primary cortices (e.g. the primary motor and sensory cortex) and relatively high ITV values in the association cortices, especially for cortical areas involved in higher-order cognitive functions, e.g. the middle frontal gyrus (MFG) and inferior parietal lobule (IPL). These high-order association areas also exhibited greater functional variability than the other parts of the cerebral cortex^17^. Similar organizational patterns were observed in the variability maps of brain morphology (modulated surface area) and connectomes (connectivity fingerprints), both of which revealed significant associations with ITV (*r* = 0.569, *p* < 0.001 for connectivity fingerprints in ***Fig. 3e***; *r* = 0.810, *p* < 0.001 for modulated surface area in ***Fig. 3f***), and regulated the topography variability through direct and indirect effects (***Fig. 3d***). Our results indicated that BAI-Net cartography captured reliable variations in topographic organization of individual brains that mainly driven by both brain morphology and human connectome. Such topographic variability was not noticeable in the conventional registration-based approach that only relies on the shape and intensity of brain structures.

**Fig. 3.**
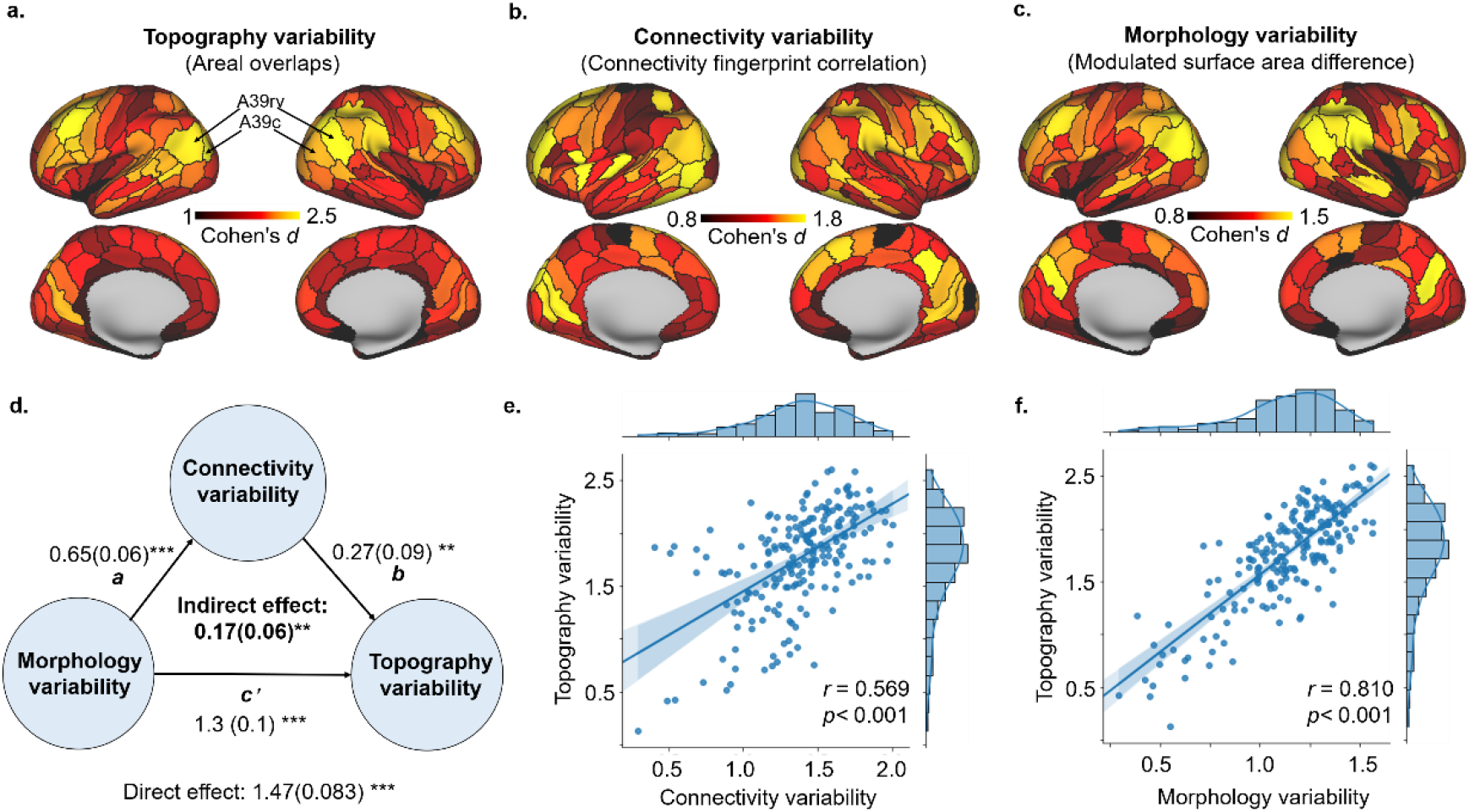
Topography variability of BAI-Net cartography was associated with and regulated by brain morphology and connectivity fingerprints. a: The distribution of inter-subject areal topography variability. b: The distribution of variability in anatomical connectivity fingerprints. c: The distribution of variability in brain morphology. d: The mediation analysis of ITV and the variability in brain morphology and connectivity. e, f: The association analysis between ITV and the variability in brain morphology (*r*=0.810, *p* < 0.001) and connectivity (*r* = 0.569, *p* < 0.001). Each dot in the scatter plot represents one cortical area.

#### Integrating area-specific connectivity fingerprints

We uncovered a long-tail distribution of ITV for BAI-Net cartographies (***Figs. 3 and 4***), with the top 10% of topography variability located in the frontoparietal regions (yellow regions in ***Fig.4a***) while the bottom 10% located in the limbic areas (blue regions in ***Fig.4a***). Different range of context information was employed in the BAI-Net cartography of these two types of brain regions. Specifically, for frontoparietal regions, the areal probability of each vertex was mainly driven by the connectivity context within a local neighborhood and itself (i.e. positive activations at K<2) while suppressing the contributions of connectivity profiles far away (negative activations at K>3), as shown in ***Fig.4b***. The limbic areas followed a similar trend but showed significant differences in the activations at different K-orders (i.e. integration scale of context). For instance, frontoparietal regions showed higher positive activations within the local filed (p-values<0.05 when K<2) and more negative activations at distributed areas (p-value<0.01 when K=5, as shown in ***Fig.4b***). These findings indicated the proposed BAI-Net cartography integrates the connectivity context from local neighborhoods, adapts the integration rule according to area-specific characteristics and captures reliable features from the integrated context of anatomical connectivity profiles.

**Fig. 4.**
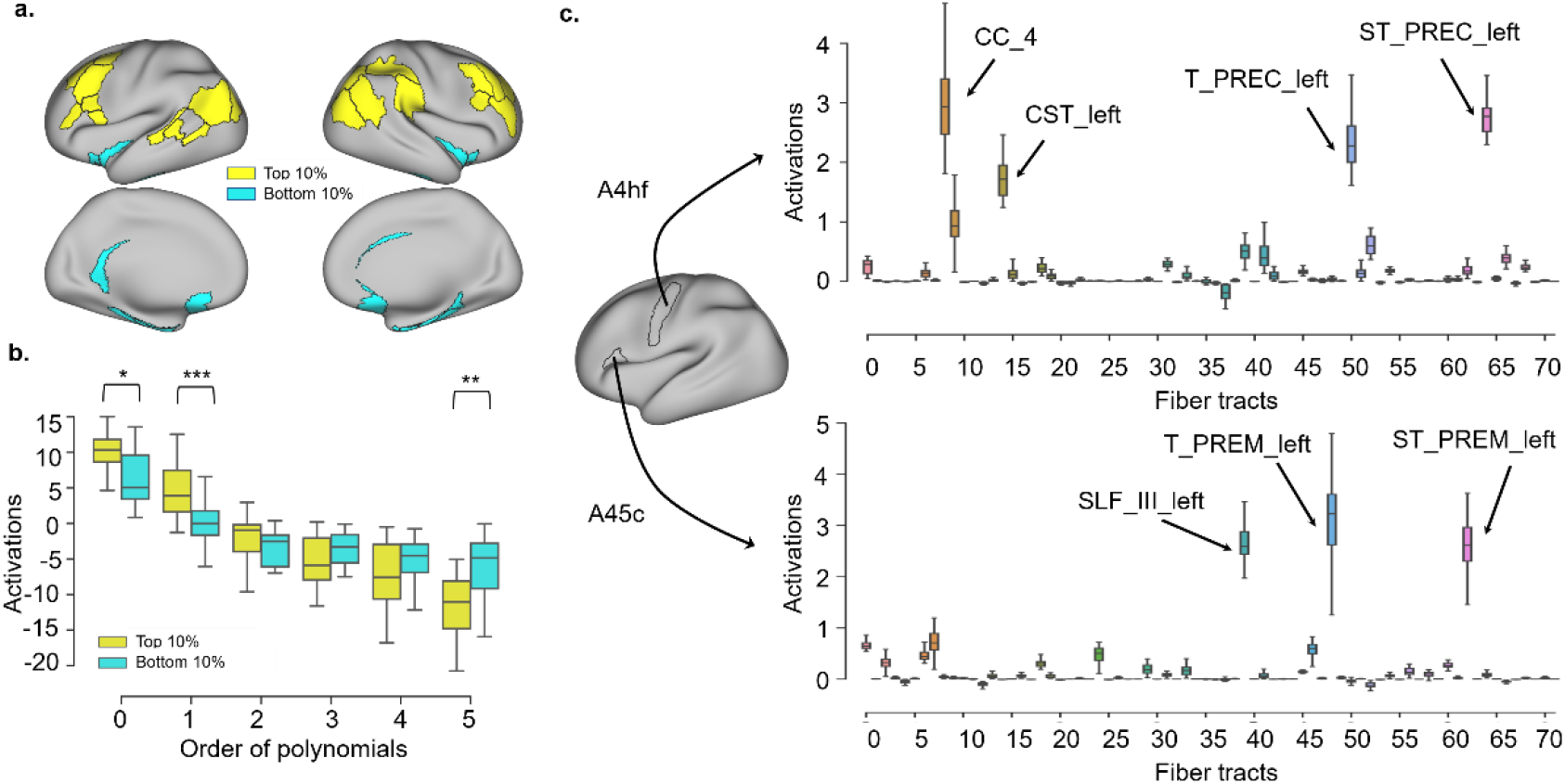
Area-specific contributing factors to the topography variability of BAI-Net cartography. a: The distribution of the top 10% and bottom 10% ITV cortical areas. b: Different activated patterns of top 10% and bottom 10% ITV cortical areas measured at different K orders in BAI-Net. c: Contributions of major fiber tracts for the areal delineation of A4hf and A45c. Different colors in the boxplot represent different major fiber tracts in the connectivity fingerprints. Note: *: *p*<0.05, **: *p*<0.01, ***: *p*<0.001.

#### Individualized cartography used biologically meaningful salient features

Whether the salient features that mostly driven the areal probability in individualized cartography are biologically meaningful is an important question. We took areas A4hf and A45c (for motor and language functions, respectively) as examples to visualize the contribution of major fiber tracts in the process of individualized cartography. As shown in ***Fig. 4c***, for areal delineation of A4hf, which was involved in the movement of the hand and face^14^, we found that the highly contributed fiber tracts included granterior midbody of corpus callosum (CC_4), corticospinal tract (CST), thalamo-precentral circuits (T PREC) and prefrontostriatal circuits (ST PREC). For areal delineation of A45c, which was involved in semantic and language processing^14, 38^, the highly contributed fiber tracts consist of the longitudinal fascicle III (SLF III), thalamo-premotor circuits (T PREM) and corticostriatal circuit (ST PREM), coinciding with the structural and connectional substrates of language processing ^38^. Our results indicated that the BAI-Net cartography captured biologically meaningful and area-specific signatures coinciding with both connectional and anatomical organization of human brain.

### Prediction of cognitive behaviors and genetic associations

#### Individualized global cartographies predicted cognitive behaviors

The topography variability of BAI-Net cartography not only significantly associated with individual variability in brain morphology and connectivity fingerprints, but also strongly predicted individual differences in cognitive behaviors. As shown in ***Fig. 5***, we trained a kernel ridge regression model for each of 58 behavioral measures and obtained 31 predictive behavioral models that showed significant predictions (*p*<0.001) using either the BAI-Net or IC approaches. The results indicated that BAI-Net method achieved higher overall prediction accuracies on 58 behavioral scores (average *r* = 0.102 and 0.078 respectively for BAI-Net and IC, paired t-test p<0.001), as well as better predictions on 31 significantly predicted behavioral measures (average *r* = 0.152 and 0.130 respectively for BAI-Net and IC, paired t-test *p*=0.028).

**Fig. 5.**
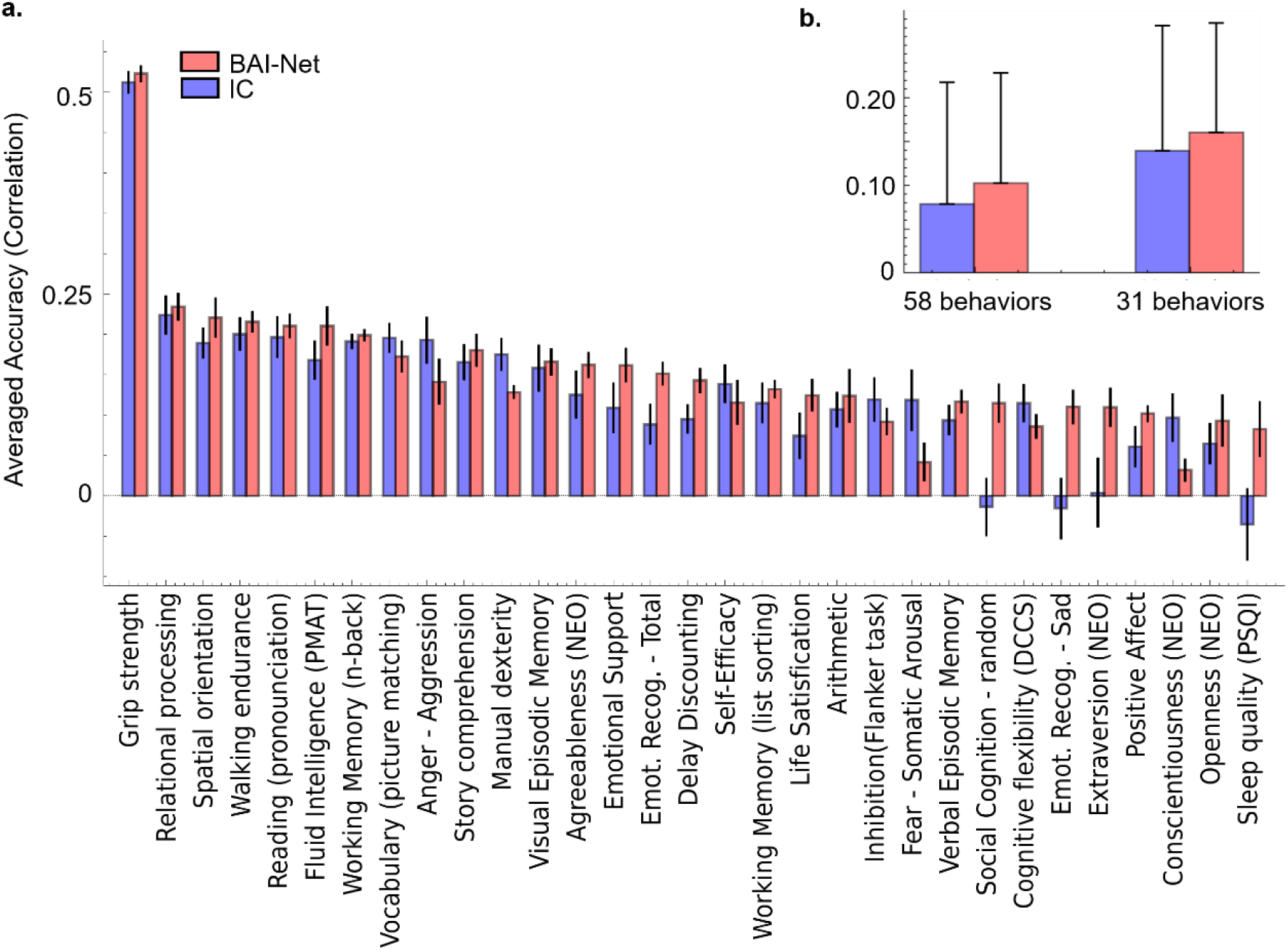
Prediction accuracy of cognitive behaviors using individualized cartographies. Using a 10-fold cross validation procedure, we evaluated the perdition accuracy on each of 58 cognitive behaviors using the topography variability from the BAI-Net or IC methods. Among which, we obtained 31 predictive behavioral models that showed significant predictions (*p*<0.001) using either the BAI-Net or IC approaches. a: Prediction accuracy on 31 significantly predicted cognitive behaviors. b: Average prediction accuracy on 58 behaviors and 31 significantly predicted behaviors.

#### Topography variability controlled by genetic effects

The topography variability of BAI-Net cartography was significantly heritable in HCP twin populations. To validate this hypothesis, we first split the whole dataset into four groups, i.e. Unrelated, Siblings, dizygotic twins (DZ) and monozygotic twins (MZ). We found that BAI-Net cartography showed more similar individualized cartographies in closed-kinship groups (***Fig. 6a***), e.g. higher similarity in the MZ than unrelated groups (Dice=0.783 and 0.733, respectively). The IC approach showed similar patterns between the four groups (e.g. Dice=0.829 and 0.816, respectively for MZ and unrelated groups) but detected smaller gaps between MZ and unrelated groups (gap=0.050 vs 0.013 respectively for BAI-Net and IC). Moreover, the topography variability of individualized cartographies was significantly impact by genetic factors, with a higher heritability value in BAI-Net (*h*^*2*^=0.175, *p*<0.001) than IC (*h*^*2*^=0.105, *p*<0.001).

**Fig.6.**
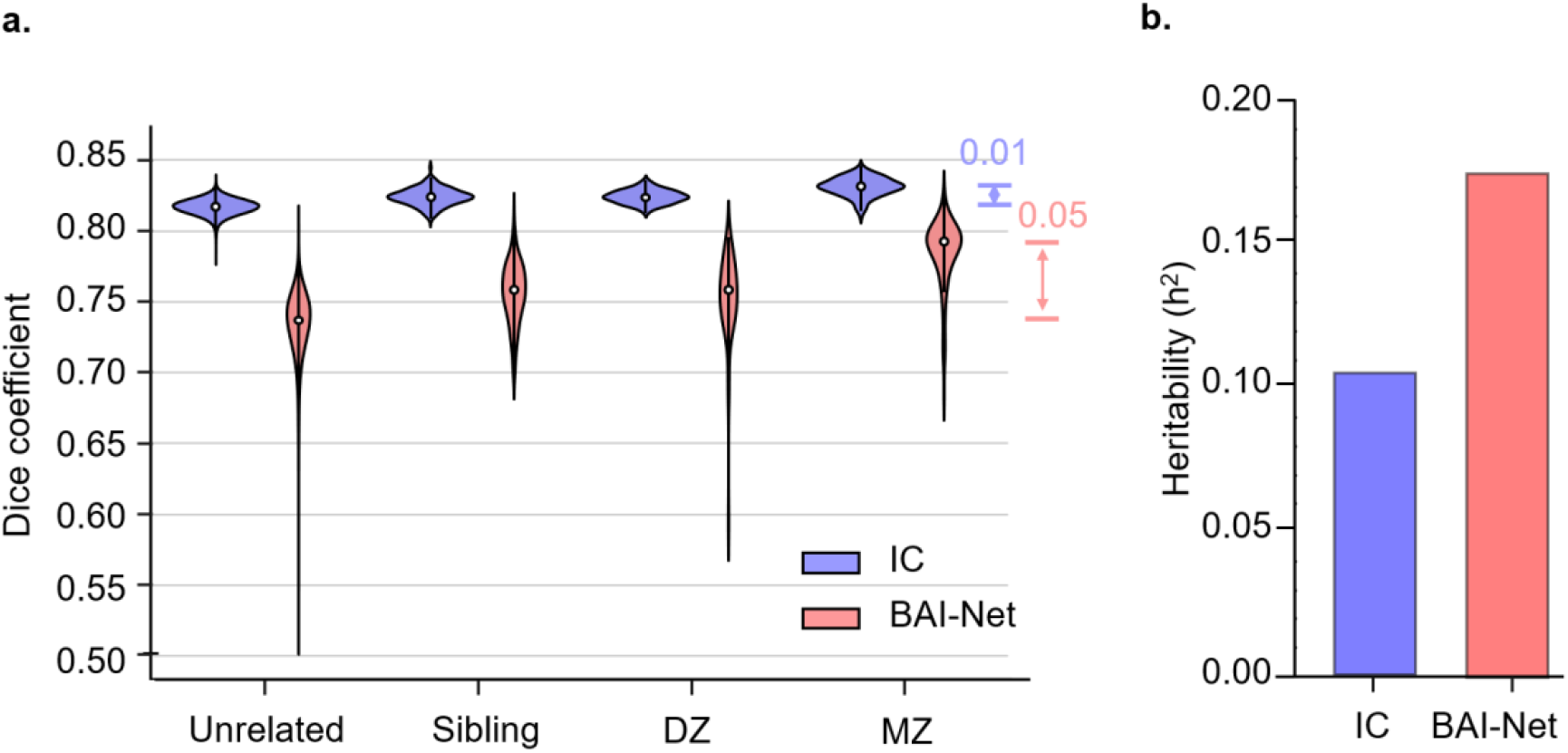
Genetic effects of the topography variability of individualized cartographies. a: Topography similarity of individualized cartographies among four different groups (Unrelated, Sibling, DZ and MZ). b: The heritability of the topography variability of individualized cartographies. The BAI-Net method showed a higher heritability value (*h*^*2*^=0.175, *p*<0.001) than the IC method (*h*^*2*^=0.105, *p*<0.001).

## Discussion

In the present study, we propose a deep-learning approach for individualized cartography which aligns the group-level cortical areas onto individual brains by taking into account the variations in brain morphology and anatomical connectomes. The proposed BAI-Net method generated highly reproducible, individual-specific cartography across various acquisition conditions, not only revealing reliable topographic patterns of a single subject across multiple sessions (Dice=0.901±0.014) but also capturing highly variable organizational principles across different subjects (Dice=0.745±0.025), yielded a significant heritability in twin population. The topography variability of BAI-Net individualized cartographies generally followed the functional and connectional gradient of the cortical organization, strongly associated with individual variability in brain morphology and connectivity fingerprints, and significantly predicted individual cognitive behaviors. Our study provides an important tool for better understanding of individual cognition, behavior, and the pathology of brain diseases and paves the way of individualized atlas-based precision medicine in clinical practice.

One of the big challenges in diffusion tractography is that massive short-distance fibers usually dominate long-distance fibers in the tractograms^30, 39^, which can easily bias the areal delineation on individual brains based on diffusion tractography maps. To overcome this issue, we used a fiber-tract embedding approach to project high-dimensional diffusion tractography maps (50k+ voxels) to a low-dimensional fiber-tract space (72 fiber bundles). This embedding technique highly reduced the computational complexity in GNN. In addition, the fiber-tract embedding focused more on the long-distance anatomical connections, largely suppressed the error accumulation effects on long-distance fibers in individual anatomical connectomes, improved the alignment of connectivity fingerprints across subjects^32, 40^, and boosted the reproducibility of individual-specific areal delineation (***Fig.2***). The majority of existing individualized cartography methods aimed for a high local homogeneity of brain signals or connectivity profiles within individualized brain parcels ^18 26^. This might not be an appropriate goal for the anatomical connectomes derived from diffusion tractography due to the distance effects. In contrast, the proposed BAI-Net method yields smaller areal homogeneity of anatomical connectivity in the tractography than the IC method (***Supplementary Results SR1***), but revealed higher variations in individual topography along with better heritability than IC(***Fig.6***).

Another challenge in the mapping of individual anatomical connectomes is the reliability and reproducibility of probabilistic tractography results ^28^. To solve this issue, we used a rich set of graph convolutions to integrate the local context of anatomical connectivity fingerprints. The local contiguity was specified by a large-scale vertex-level graph (32k vertices per hemisphere) with each node indicating a cortical vertex and each edge indicating whether two nodes shared in a triangle in individual cortical surfaces. The usage of such brain graph implicitly applied a smoothing effect on the cortical surface such that adjacent vertices on the graph had similar connectivity fingerprints and graph representations. Besides, in contrast to previous individualized parcellation approaches which only used the connectivity information of the target vertex (e.g. the IC method ^26^), our model also took into account the local context of connectivity fingerprints within a specified neighborhood. The contributions at different spatial neighborhoods (K-orders) were dependent on the nature the brain parcels and varied a lot across different brain regions (***Fig.4***). Together, the deep graph convolutional architecture combined the vertex graph and connectivity context in the model and ensured the balance between individual-specific topography and populational aligned organizational principles. The BAI-Net model generated highly reproducible and robust individualized cartographies, not only aligning with the global topographic pattern specified in a group atlas but also revealing considerable differences in individual topography that strongly associated with the anatomical gradients in both brain morphology (*r*=0.810) and anatomical connectivity (*r*=0.569) (***Fig.3***). Lastly, the usage of group-prior constraints further enhanced the inter-subject alignment in individualized cartographies and might ensure the significant predictions of individual cognitive behaviors (***Fig.5***) as well as its consistency on repeated sessions and in the twin populations.

Individualized brain cartography has played a more and more important part in neuroscience research and clinical studies. Accumulating evidences suggested considerable and meaningful variations in individual topography at the levels of brain areas and networks. ^18-20, 26, 33, 41^. Despite various imaging features used in individualized cartography methods so far, we still observed high consistency in terms of the variability in individual topography (ITV) by using either anatomical or functional connectomes. First, the anatomical ITV revealed by BAI-Net exhibited small values in the primary cortices (e.g. the primary motor and sensory cortex) and relatively high ITV values in the association cortices, especially for cortical areas involved in higher-order cognitive functions including MFG and IPL, coinciding with the functional ITV maps despite of using different group atlases ^20^.More importantly, the anatomical ITV showed similar predictability on individual cognitive behaviors (*r*=0.102 for 58 cognitive behaviors) as compared to the functional ITV (*r*=0.111 for 58 cognitive behaviors) ^20^, and even achieved higher prediction accuracies on some behavioral measures (e.g. grip strength and fluid intelligence shown in ***Fig.6***). Besides, the anatomical ITV exhibited much higher reproducibility (Dice=0.90) as compared to the functional ITV (Dice=0.81 for 400 cortical areas ^20^, Dice=0.78 for 17 networks^19^)

The reproducibility of individualized cartography is an essential requirement in clinic practices, long with the robustness, interpretability and time-consumption. The BAI-Net method achieved high specificity (Dice=0.703), high robustness (Dice=0.83) across various acquisition conditions on different scanners. Then area-specific activated rules in spatial activations interpretates the areal identification process of BAI-Net method. Meaningful and salient connectivity fingerprints was captured and contributed more in areal identification. For example, the SLF III, T PREM, ST PREM elements of the connectivity fingerprints contributed more in A45c area, which coincides with the structural and connectional substrates of language processing^14, 38^. More importantly, with a preprocessed individual cortex surface, the time consumption of the BAI-Net method to inference the cerebral cartography of a new subject was around 2∼3h on the Centos 6 Linux system from raw diffusion images. But it will cost less time in the regional cartography of BAI-Net method according to the size of the region, which might partially solve the time problem of probabilistic tracking when regional areas are needed. As mentioned above, the BAI-Net satisfies these necessities for the clinic applications.

The BAI-Net method provides a generalized framework for individuated cartography with the assistance of graph neural networks. The presented method is not limited to a certain group atlas, a similar implementation of BAI-Net using Glasser’s atlas ^13^ was shown in ***Supplementary Fig. S10***. In clinical applications, a faster, reliable, individual-specific mapping of the cerebral cortex is the critical step towards for personalized precision medicine, which enables the personalized localization of neuroimaging biomarkers, the investigation of individualized structural-functional relations, and potentially assist the development of new technologies in practical treatments, e. g. locating the target areas for transcranial magnetic stimulation (TMS) and deep brain stimulation (DBS) therapies, and reducing functional impairment in neurosurgery.

## Materials and Methods

### Datasets and preprocessing

#### Dataset 1: Human Connectome Project (HCP)

We acquired healthy young adults from the HCP S1200 release, consisting of T1 weighted (T1w) data, resting-state functional MRI (rs-fMRI), as well as diffusion MRI (dMRI) data for each subject. Specifically, the BAI-Net method was trained and evaluated on the first dataset, consisting of 969 subjects (Age: 21-35, Female: 519) acquired from the Human Connectome Project S1200 release (after removing subjects with large head motions). The test-retest reliability of the model was then evaluated on the second dataset, consisting of 44 subjects acquired from the HCP test-retest datasets. The preprocessed datasets were used in the current study using the HCP minimal preprocessing pipeline ^42-44^. The individual cortical surfaces were first reconstructed from T1-weighted MRI data and then projected onto the standard surface template (fs_LR_32k) with 32k vertices per hemisphere by using the MSMAll registration approach ^23^. Diffusion MRI data had been mainly preprocessed by motion correction, eddy current distortion correction, and echo-planar images (EPI) susceptibility-induced field distortion correction. The preprocessing of the functional MRI data mainly included motion correction, EPI susceptibility-induced distortion correction, linear trend removal, and independent component analysis (ICA)-based artifact removal. Further preprocessing details can be found in the HCP preprocessing pipeline (https://github.com/Washington-University/HCPpipelines) ^45, 46^. In heritability analysis, all HCP subjects were divided into for four groups, 1) Unrelated (468119 pairs): no shared parent IDs; 2) Siblings (297 pairs): sharing one parent ID and but not twins. 3) DZ (60 pairs): dizygotic twins according to the genetic records. 4) MZ (119 pairs): monozygotic twins according to the genetic records.

#### Dataset 2: MASiVar dataset

Additional dataset was used acquired from the Multisite, Multiscanner, and Multisubject Acquisitions for Studying Variability (MASiVar) dataset ^47^, consisting of 74 scans (removed 8 scans due to incomplete brain tissues in diffusion images) and 14 healthy adults (8 males and 6 females, age 27-47). This dataset was used to evaluate the stability of individualized cartography on multiple sessions, sites, and scanners. Specifically, dMRI data was acquired from 3 cohorts using four different scanners (two 3T Philips Achieva scanners at two different sites, one 3T General Electric Discovery MR750 scanner, and one 3T Siemens Skyra scanner) with at least one T1-weighted image for each subject at each session. Of the three cohorts, different scanning sequences were used in diffusion imaging, including different b-values (b = 1000 to 3000 s/mm^2^) and diffusion directions (about 40∼96 directions for each b value), different spatial resolutions ranging from 2.5 mm isotropic or 2.1 mm by 2.1 mm by 2.2 mm (sagittal, coronal, and axial), and so on. These diffusion images were preprocessed using the PreQual pipeline ^48^ with the default settings, including intensity normalization and distortion correction, as well as the Marchenko-Pastur technique^49-51^. More information on PreQual can be found at https://github.com/MASILab/PreQual.

### Pipeline of BAI-Net individualized cartography

The Brain Atlas Individualization Network (BAI-Net) pipeline included three key steps: construction of individual cortical graph, embedding of individual connectivity fingerprints and node classification using the connectivity context.

#### Step1: Construction of individual cortical graph

The graph structure of individual brains was derived from the cortical mesh data of each subject. For each hemisphere of each subject, we constructed a spatial brain graph, with nodes indicating each cortical vertex (consisting of around 30k vertices/nodes after excluding confounding vertices in the medial wall), and edges indicating whether two vertices shared in a triangle of the cortical surface, weighted by the inverse of the geometric distance between them. The brain graph was sparsely connected and highly localized in space, with each vertex connecting with 2-6 nearest vertices on average. The brain graph architecture provided a reference structure for each vertex to search for its spatial connectivity context.

#### Step2: Embedding of individual connectivity fingerprints as node features in the graph

After constructing the graph, the connectivity fingerprint of each vertex was calculated as follows: 1) A surface-based probabilistic tractography algorithm was applied to track 5000 streamlines from each vertex on the cortical surface throughout the whole brain, including both cortical and subcortical regions ^52-55^. The resulting whole-brain tractography map 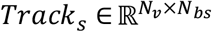 (*N*_*v*_ is the number of surface vertices and *N*_*bs*_ is the numbers of brain voxels of subject *s*), was first threshold at 2 and then down-sampled to a 3mm-resolution space ^56^. 2) A binary mask was created for each of 72 fiber bundles (names of fiber bundles listed in ***Supplementary Table S1***) by using a pretrained deep-learning model named TractSeg ^31^. The resulting fiber-tract mask was also down-sampled to 3mm resolution. 3) An embedding of the whole-brain tractography map was created for the projection from the volume space (50k+ voxels) to the fiber-tract space (72 fiber bundles). After that, the connectivity fingerprint 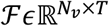 and (*N*_*v*_ ≈ 30k and *T* =72) was generated as follows:

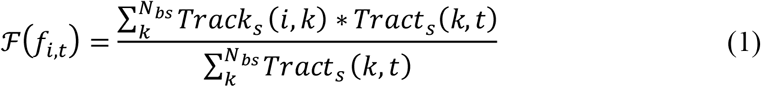

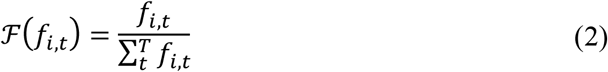

where each element *f*_*i,t*_ in ℱ indicates the probability of any fiber-tract *t* existing in the tractography map of vertex *i*. These connectivity fingerprints are biologically meaningful and have been used to locate similar functional areas across different species^32^.

#### Step3: Node classification using the connectivity context

Most of current cortical parcellation strategies predicted the area/parcel assignment solely based on the connectivity information of the target vertex while neglecting the context information of the connectivity profiles ^57^. The connectivity context starts to show potentials in the field of brain cartography. Cohen and his colleagues proposed to detect local changes in functional connectivity maps through an edge detection algorithm on the cortical surface ^58^, which only used the local context from the first-order neighborhood (directly connected vertices). Graph neural networks provide a more generalized approach for integrating the context information at each node. One type of graph convolution of *x* using Chebyshev polynomials is defined as:

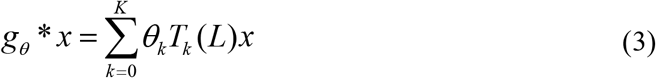

where 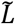 is a normalized version of the graph Laplacian and is equal to 2*L*/*λ*_*max*_ − *I*, with *λ* being the largest eigenvalue. *θ*_*K*_ is the parameter to be learned for each order of the Chebyshev polynomial *T*_*k*_, (*T*_*K*_(*x*) = 2*T*_*K*−1_(*x*) − *T*_*K*−2_(*x*), *T*_0_(*x*) = 1, *T*_1_(*x*) = *x*). By using a truncated expansion of the Chebyshev polynomial (as shown in Eq.1), the ChebNet graph convolution is naturally K-localized in the graph ^59^. Specifically, when *K*=1, the graph convolution only considers the context information from the direct neighbors at each node. When *K*>1, the graph convolution also takes into account the information from a larger scale neighborhood, including nodes that can be reached within K steps of random walk on the graph. All this information is then integrated using the graph convolution. A stacked two-layer GNN architecture was used to enlarge the receptive fields of information integration. The GNN model took individual brain graphs as inputs, integrated the context information of brain connectivity at each vertex, and generated new representations (shown in ***Fig. 1b***).

### Optimization of BAI-Net method

The constructed individual brain graphs were used to train a GNN model constrained by group priors (***Fig. 1b***). Specifically, an example of the group prior was extracted from 210 areas on the cortical surface of the Human Brainnetome Atlas, defined based on a group of 40 healthy subjects. The group atlas was first mapped onto the standard surface template (fs_LR_32k) using the MSMAll approach ^23 60^. A one-hot encoding of the areal label was created for each vertex (in total of *C+1* dimensions, where *C* is the number of parcels in the group atlas), and then used to modify the loss function for the GNN. Specifically, the GNN model takes an individual brain graph 𝒢 = (𝒱,*ε*,ℱ) as inputs, where 𝒱 is the set of vertices (around 30k per hemisphere) in the cortical surface, *ε* is the set of edges, showing whether two vertices share a common triangle in the surface, and 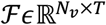 is the set of feature vectors indicating the connectivity fingerprints *f*_*i*_ defined on each vertex, *N*_*v*_ is the number of vertices, *T* is the number of fiber tracts (here *N*_*v*_ around 30k and *T* =72). Using the validation dataset, optimal parameters for the two-layer GNN model is chosen. The learned graph representations extracted from the last layer were then projected to a (C+1)-dimensional probability vector at each vertex using the SoftMax function. The K-L divergence was used to calculated the difference between the group prior *y*_*i,k*_ (one-hot encoding) and the predicted areal probability *p*_*i,k*_ at each vertex *v*_*i*_ for each region *k*. The weight of uncertainty *w*_*i*_ at each vertex was inferred from the populational probability map of the group atlas. Thus, the final loss function was defined as follows:

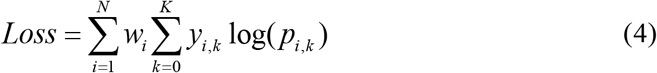

The benefits of using the above loss function include: 1) high contributions of the vertices near the center of regions help to obtain a high level of inter-subject alignment for the global topographic organization. 2) small contributions of the vertices at the borders of regions help retain the inter-subject variability to some degree and allow the mismatching of label assignments in the local architecture.

The hyperparameters of the model was determined in the validation set. A 5th-order graph convolution was used in GNN with *M=32* convolutional kernels in the first GNN layer and *C+1=106* kernels in the second GNN layer. We used Adam as the optimizer with an initial learning rate of 0.01 (decreased to 0.0001 after the 5th epoch). An additional L2 regularization of 0.0005 on weights was used to control model overfitting and noise in the imaging data. The network was trained on 100 unrelated subjects for 50 epochs with the batch size set to 1 (processing one subject at a time on the 12G GeForce GTX 1080K) using traditional dataset and random-order dataset respectively and evaluated on the validation set of 20 subjects at the end of each training epoch. The best model over 50 training epochs, that is, the one that achieved the lowest loss on the validation set, was saved and further evaluated on the independent test set. During the model evaluation (***Fig. 1c***), an individual brain graph was first constructed from the test subject, using surface construction, diffusion tractography, and fiber-tract embedding. Then, the GNN model took the brain graph constructed from the target subject as input, predicted the areal probability vector at each vertex and labelled graph nodes with the highest probability.

### Group-registered map and iterative clustering

We included two approaches as the baseline approaches in this study. First, the group-registered map was generated by mapping the original group atlas from the MNI volume space to high-resolution surface template (164k), and down-sampling to the fs_LR_32k surface template, and then projecting onto individual surfaces.

The second approach was to iteratively assign each vertex to different brain parcels and update the connectivity information of each area until model convergence. This iterative clustering approach was originally proposed for fMRI-based individualized brain cartography, we adapted it onto diffusion tractography based individual cartography (detail descriptions can be seen in ***Supplementary Methods*)**.

### Robustness, specificity and inter-subject variability

The robustness and specificity of individual cartographies were evaluated by the areal overlaps (Dice coefficient) between intra-subject and inter-subject pairs, using HCP test-retest dataset as well as multi-scanner MASiVar dataset. In the calculation of areal overlaps, the surface cartography for each subject was first converted into binary ROIs (C cortical areas in each hemisphere, one for each area), and were concatenated into a single vector. The Dice coefficient was calculated with the equation (2*A∩B)/(A + B) between two vectors ^13^ for any area overlap (topography similarity) mentioned in this article. All cortical surfaces were created using the Connectome workbench toolbox.

Inter-subject variability of a property was estimated by the effect size, Cohen’s *d*, which revealed the real inter-subject variations after removing the intra-subject variations. Thus, inter-subject variability (the effect size) was defined as:

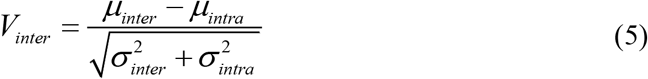

Where *µ*_*inter*_ and *σ*_*inter*_ represented the mean and standard deviation of the variabilities between each pair of subjects, while *µ*_*intra*_ and *σ*_*intra*_ represented those in the variabilities between different scans for the same subject.

Different measures were used to calculate inter-subject variabilities, including areal topography (areal overlaps), areal morphology (modulated surface area), and areal connectivity (connectivity fingerprint). Modulated surface area was calculated by the averaged vertex area within a cortical region (‘surface-vertex-areas’ command in Connectome Workbench toolbox). The variability in brain connectivity. The variability in areal topography was measured by 1 – *Dice* (topography similarity of two individual cortical areas). The variability in brain morphology was measured by the absolute difference of two modulated surface area. The variability in brain connectivity was calculated by 1 – *r* (Pearson correlation of two area-averaged fingerprints). Additional correlation analysis was performed between the inter-subject variability of areal topography, morphology and connectivity.

### Model activations in the BAI-Net model

The model activations in the 1st GNN layer of BAI-Net were analyzed and interpreted in two aspects: activations at different spatial ranges and contributions of each connectivity fingerprint. The order of the Chebyshev polynomials (K-order) can be regarded as the distance from each cortical vertex to the related connectivity context, ranging from *K = 0* (the vertex itself) to *K = 5* (connected to the vertex through five steps on the graph).The activations at different K-orders were averaged across all the graph filters with positive activations indicating supporting the identification of the target area and negative activations indicating suppressing the contributions of connectivity fingerprints to the target area. For the delineation of each area, the contributions of each major fiber tract was estimated in two steps: 1) selecting all the positively activated graph filters from the model; 2) calculating the averaged activation of each fiber tract in the selected graph filters. The activations of the major fiber tracts were regarded as the contributions of connectivity fingerprints.

### Prediction of individual cognitive behaviors

The topography variability of individual cartographies could reflect the individual idiosyncrasies^20^. Here, we adopted the kernel ridge regression with L2-norm regularization to predict the individual cognitive behaviors. We used the Dice coefficient as the kernel function *K*(·) and regularization parameter *α=1* in the prediction model:

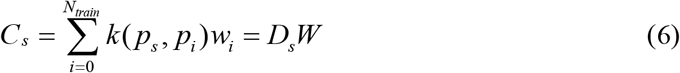

where 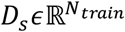 indicates the topographic similarity (measured by Dice score) between the selected cartography *p*_*s*_ with the training cartographies *p*_*i*_, *i* = 1, …, *N*_*train*_ ; *C*_*s*_ indicate the predicted behavioral scores for subject *s*; *W* indicates the regression parameters on training subjects. The objective function was defined as follows:

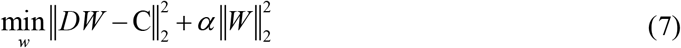

We trained a prediction model for each behavioral score and evaluated the model using a 10-fold cross-validation procedure. The prediction accuracy on the test fold was evaluated by the Pearson correlation between all predicted and actual behavioral scores. The averaged accuracy on the 10 folds was reported as the final performance.

### The heritability of individual-specific topography

The topography of individual functional networks can be explained proportionally by the genetic variation among individual in a population ^61^. The genetic effect of the topography should also be revealed in the region-level cerebral cartography. So, we calculated the heritability of individual-specific topography according to the scripts in https://github.com/kevmanderson/h2_multi/blob/master/h2_multi/h2_multi.m. A multivariate linear mixed effects model was built as follows:

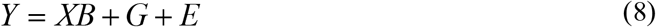

where *Y* was the multi-dimension phenotype, *X* was the covariates (including age, sex, age^2^, sex×age, age^2^×sex, total surface area and FreeSurfer-derived intracranial volume.), *B* was the fixed effects, *G* was the additive genetic effects from single nucleotide polymorphism (SNP) and *E* was the unique environmental factors. The detailed calculation of the heritability can be seen in the article ^61^.

## Data Availability

The pipeline of BAI-Net method will be open-sourced on Github website once accepted. And other data and figures of this article are available on request from the authors.

## Author contributions

The Liang Ma and Prof. Yu Zhang contributes to the writing, coding, plotting and validation of the pipeline. And Hantian Zhang, Luqi Cheng, Yuheng Lu provides ideas and codes in the individual pipeline. Others provides several suggestions and corrections in the writing and validation of the pipeline. Prof. Tianzi Jiang and Prof. Lingzhong Fan contributes equally in the support of the publication of this article.

## Acknowledgements

This work was partially supported by the Natural Science Foundation of China (Grant Nos. 91432302, 31620103905 and 81701781), the Science Frontier Program of the Chinese Academy of Sciences (Grant No. QYZDJ-SSW-SMC019), National Key R&D Program of China (Grant No. 2017YFA0105203), Beijing Municipal Science & Technology Commission (Grant Nos. Z161100000216152, Z161100000216139, Z171100000117002), Beijing Advanced Discipline Fund, the Guangdong Pearl River Talents Plan (2016ZT06S220), Major Scientific Project of Zhejiang Lab (No. 2020ND8AD02 and No.2021ND0PI01), Youth Innovation Promotion Association Chinese Academy System (CAS).

