## Supplementary material for "BAI-Net: Individualized Anatomical Cerebral Cartography using Graph Neural Network": supplymentary method, materials and figures

### Supplementary Information for BAI-Net: Individualized Anatomical Human Cerebral Cartography using Graph Convolutional Network

#### Supplementary Methods

##### The surface-based iteration clustering method for anatomical connectivity

Original volume-based method can be seen in Han's article[1]. A proper adjustment on surface is made, the details surface parcellation is conducted below:

Step 1: Probabilistic tractography was conducted in the same manner like BAI-Net methods, in which used cortical surface vertices as seeds and made whole-brain tracking. The whole-brain tractography was then down-sampling to 3mm brain mask as the final anatomical connectivity profile of each vertex. The 105 reference connectivity profiles were calculated initially by averaged connectivity profiles in the surface location that registered by group atlas.

Step 2: The connectivity profile resulting from probabilistic tractography of each surface vertices in the individual subject was correlated to the 105 reference connectivity profiles. Each vertex was assigned to 1 of the 105 regions based on maximal correlation with the reference connectivity profiles. In addition, a confidence value is calculated as the ratio between the maximum and the second largest correlation values.

Step 3: New reference connectivity profiles were generated, and then each vertex in the cortical surface was reassigned. After all vertices were assigned to 1 of the 105 subregions in the previous step, a mean connectivity profile was calculated in each cortical region, during which the diffusion connectivity profiles of the vertices with a confidence value greater than a preselected threshold (here it was set to 1.1) were averaged with greater weight. This weighting strategy guaranteed that the connectivity profiles coming from core vertices with higher confidence levels were weighted more than connectivity profiles of the vertices located at the edge of cortical regions. Then, for each region, the mean connectivity profile and the reference connectivity profile were averaged in a weighted manner again, during which the reference connectivity profiles were weighted more than the mean connectivity profiles to slowdown the convergence speed. The resulting connectivity profile estimate was utilized as the new reference connectivity profile for the next iteration. Then, the cortical vertices were further reassigned to 1 of the 105 regions using these new reference connectivity profiles.

Step 4: The iteration procedure reached convergence. Step 3 was iterated until the algorithm reached a predefined stopping criterion. Here, the iteration procedure was stopped when the parcellation pattern remained the same for 98% of the vertices in 2 consecutive iterations.

##### Statistic scores of individual cortical areas

We evaluated the performance of areal features using below statistic scores using the testing dataset. The statistic scores were listed below:

**Areal detection rate:** we also considered the areal sizes of the resulting subject parcellations relative to the population priors. If an area in an individual model was within 0.33x to 3x the size of the group area, this area was considered as being detected for this subject.

**Areal overlaps with priors:** the overlaps between each individual parcel and corresponding group locations were calculated. The areal overlaps were averaged for all areas across subjects.

**Similarity of maximum probability map**: the maximum probability maps (MPM) were annotated as the area with the highest probability at each vertex. The similarity is calculated using areal overlaps between the MPM of individual parcellations and the group atlas.

##### Statistic scores of areal anatomical connectivity

The anatomical connectivity scores of an individual parcellation were investigated from two aspects: the homogeneity of anatomical connectivity and the inter-subject similarity of connectivity fingerprint:

**The homogeneity of anatomical connectivity**: it was calculated by the average Pearson correlation coefficient between each pair of whole-brain anatomical connectivity between vertices within a cortical area.

**The inter-subject similarity of connectivity fingerprint:** the connectivity fingerprint similarity of a cortical area was first calculated by the averaged Pearson correlation coefficient of areal connectivity fingerprint across subjects. Then the similarity of all cortical areas was averaged, weighted by the vertex number of each area. We tested the significance of population scores between two methods using the bootstrap technique (1000 samplings, 30 subjects per sampling).

### Supplementary Results

##### SR1: BAI-Net method focuses on cross-subject similarity of anatomical connectivity

We investigated the anatomical connectivity from two aspects: the homogeneity of anatomical connectivity and inter-subject similarity of connectivity fingerprint. focus on the consistency of anatomical connectivity within a single subject while the inter-subject similarity of connectivity fingerprint focus on the consistency of anatomical connectivity across subjects. As a result, it has shown an improvement in the homogeneity of anatomical connectivity for both IC and BAI-Net methods. As shown in ***Fig S6***, the BAI-Net methods increased on average to 0.256, (paired $T$ test, n=50, with $p < 0.001$), compared to the registration-based method (0.243). While the IC method have shown a better improvement with 0.267 than BAI-Net method (paired $T$ test, n=50, with $p < 0.001$). However, for the inter-subject similarity of connectivity fingerprint, it shown higher consistency across subjects using the BAI-Net method than the IC method on average (BAI-Net: 0.970; IC: 0.957; two-sample t test, p<0.001). A t-SNE visualized mapping of areal fingerprints in subareas in ITG regions across subjects was shown in ***Fig. S7***. The points from the same areas is more clustered and concentrated in BAI-Net method than IC and group-registered methods.

The two measurements reflected the difference focuses of two methods. The IC method focused on individual information more, despite it had group connectivity priors which was calculated on group locations. While BAI-Net method focused more on cross-subject alignment. In other words, the areas identified by BAI-Net method might be more suitable for cross-subject comparison, because the anatomical connectivity of these areas was more consistency across subjects. It gave a substantial basis in the cross-subject analysis for the individual-specific topography for the prediction of individual behaviors and the heritability analysis (shown in ***Fig. 5*** and ***Fig. 6***).

#### SR2: Regionally improvement of the homogeneity of functional connectivity

A comparison of functional connectivity profile in regions were conducted between BAI-Net and group-registered methods as well as between the IC and group-registered methods. We took the inferior parietal lobe (IPL) region as example, which has been parcellated into 7 subregions, each consisting of distinct functional and anatomical connectivity profiles. Here, we specifically focused on two subregions, A39rv and A39c, to show whether there is a better functional alignment. The aim of this experiment was to shown the RSFC of different delineation region of A39rv and A39c among two methods was more similar with the RSFC of two areas.

As shown in ***Fig. S8***, a large difference was detected between the BAI-Net and group-registered method (the black lines delineating the borders between A39c and A39rv in the group-registered method). The differential region between the two methods (in red) was separated from the other two common regions (A39rv, in brown, and A39c, in blue). Resting-state functional connectivity maps (RSFC) were calculated for all three regions of interest. Overall, the RSFC of the differential region (which has been assigned to A39rv in the BAI-Net method and A39c in the group-registered method) showed significantly higher similarity with the RSFC of A39rv (in brown, $r = 0.70$, $p<0.001$), compared to the RSFC of A39c (in blue, $r = 0.48$) across test sessions (two encoding directions * two runs) in the HCP test-retest datasets. However, with the same analysis in the IC method, the RSFC of the differential region only showed a little higher (in brown, $r = 0.59$, two-sample t test, $p=0.708$), compared to the RSFC of A39c (in blue, $r= 0.57$). This indicated that the BAI-Net method provided a better alignment of functional connectivity profiles regionally.

### Supplementary Figures

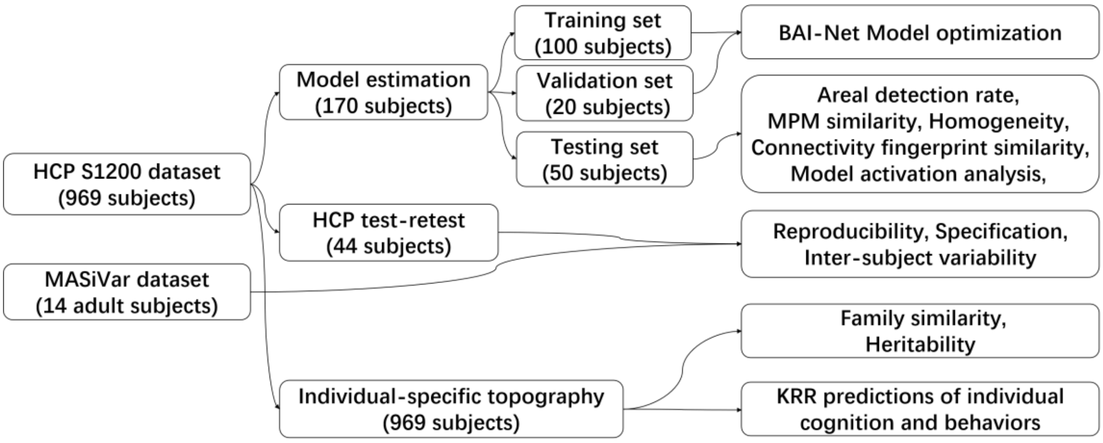


Figure S1. The evaluation datasets in different evaluation steps.

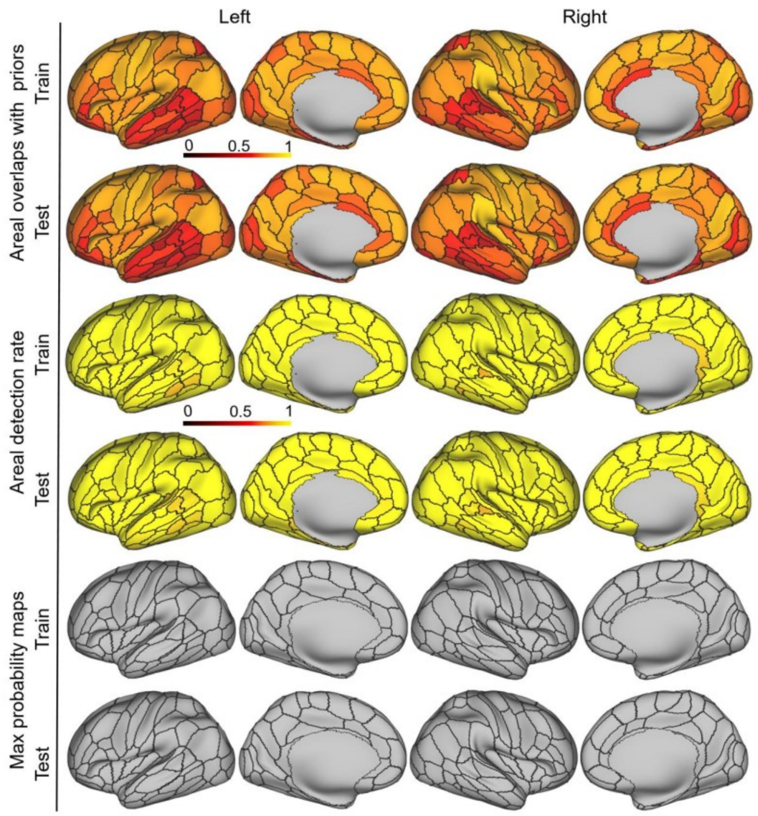


Figure S2. Reproducibility and consistency of the individual parcellations using group priors. The first/second rows (training/test): the areal overlaps of the individual parcellations with group atlas. The third/fourth rows (training/test): the areal detection rate of BAI-Net. The detected area size was between 1/3 times and 3 times the group area size. The fifth/sixth rows: Similarity of maximum probability map with group atlas.

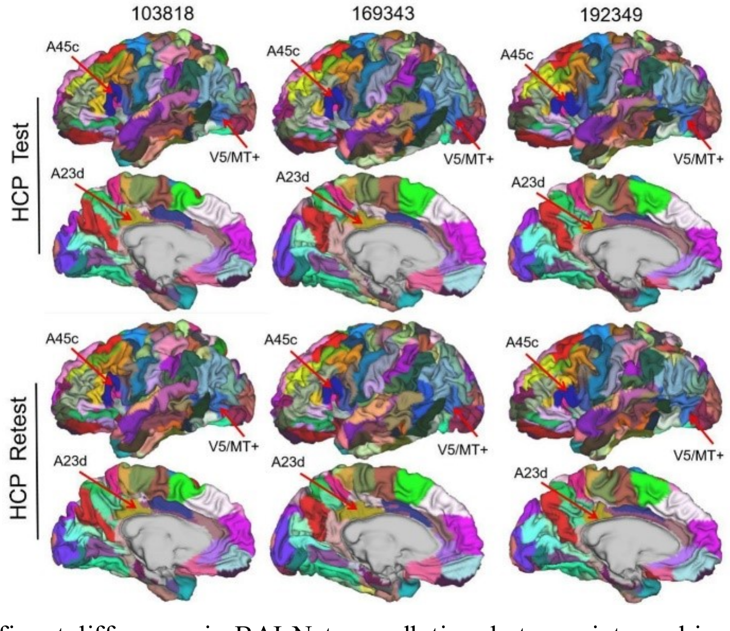


Figure S3. Significant differences in BAI-Net parcellations between intra-subject and inter-subject pairs in the left hemisphere of three randomly selected individuals

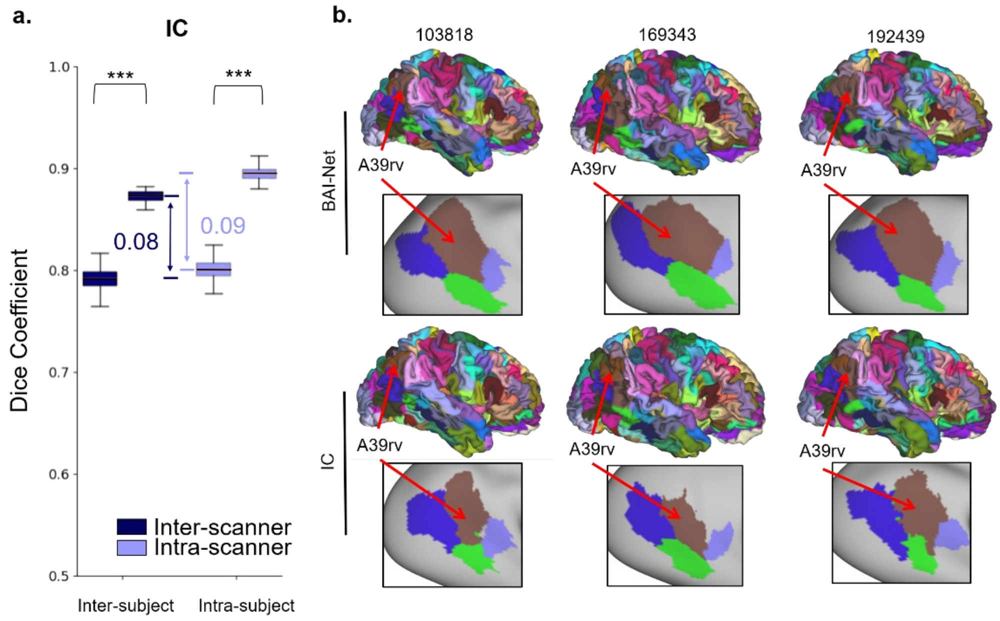


Figure S4. The reproducible and specificity of the IC method. a: the multi-scanner performance using MASiVar dataset. The differences between IC and BAI-Net methods using HCP test-retest dataset.

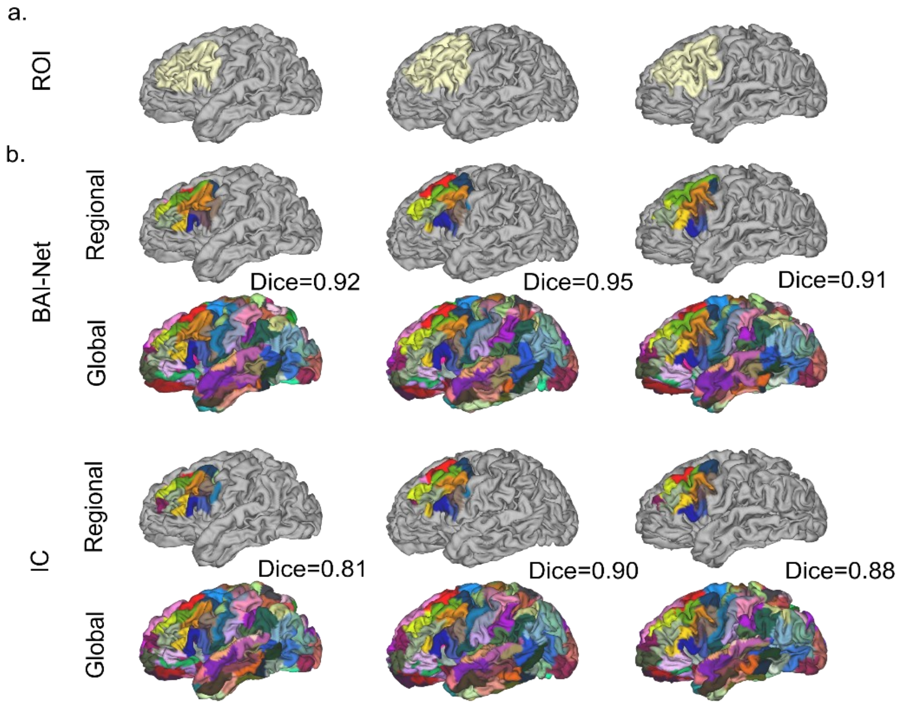


Figure S5. Regional cartography of two method for dlPFC regions. a: the ROI selection on cortical surface within a 5cm geodesic line. b: BAI-Net and IC cartographies in the areal overlaps of local and global mode using the same subjects (subject ID: 103818, 169343, 192439). There are high overlaps for the BAI-Net method than the IC method in the cartography.

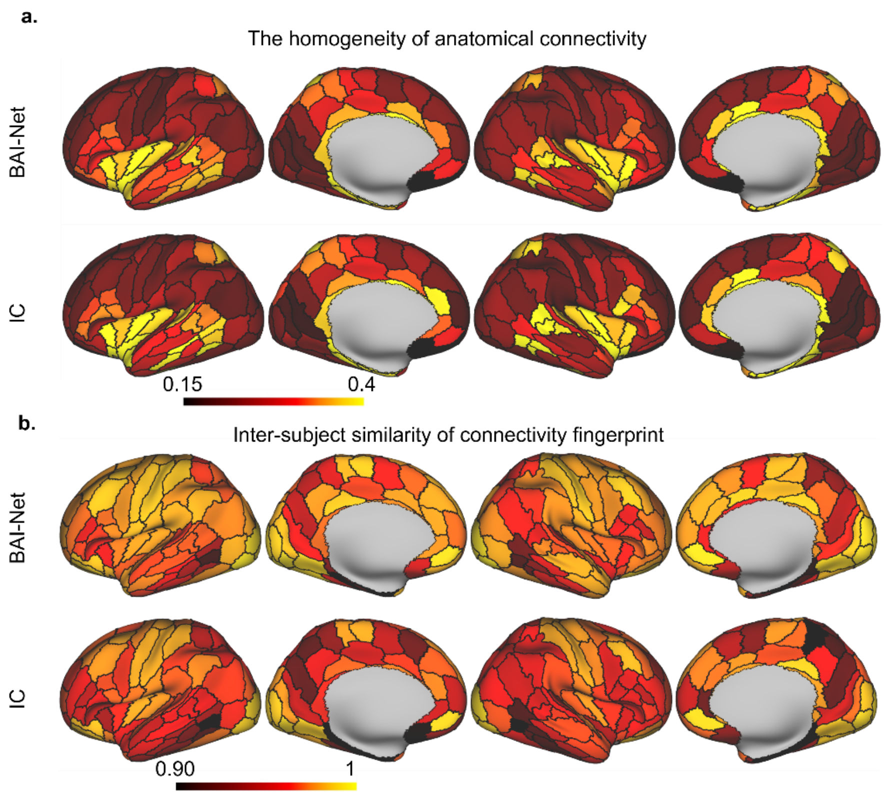


Figure S6. The anatomical connectivity scores of the IC and BAI-Net methods. a: the homogeneity of anatomical connectivity. b: inter-subject similarity of connectivity fingerprint.

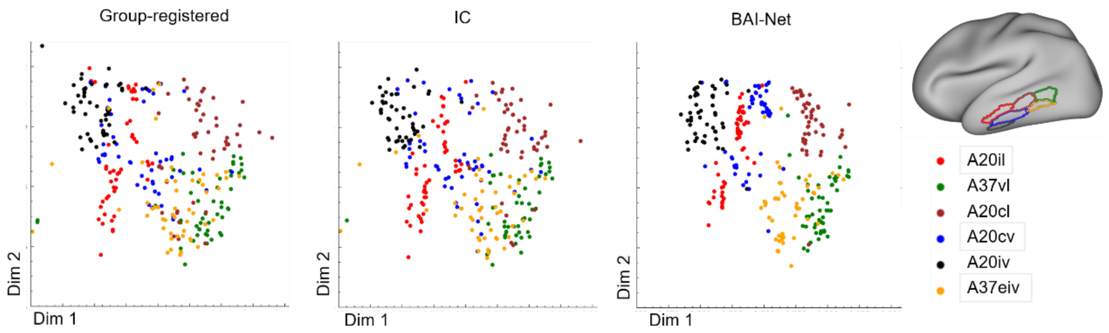


Figure S7. t-SNE projections of areal connectivity fingerprints for ITG region across subjects. The connectivity fingerprints were averaged within each subregion for ITG region. The subregions were either acquired from the group-registered, the IC and the BAI-Net method. Each dot in the plot represents one subject from the HCP test set (50 subjects in total), and each color represents one of the subregions in the target area.

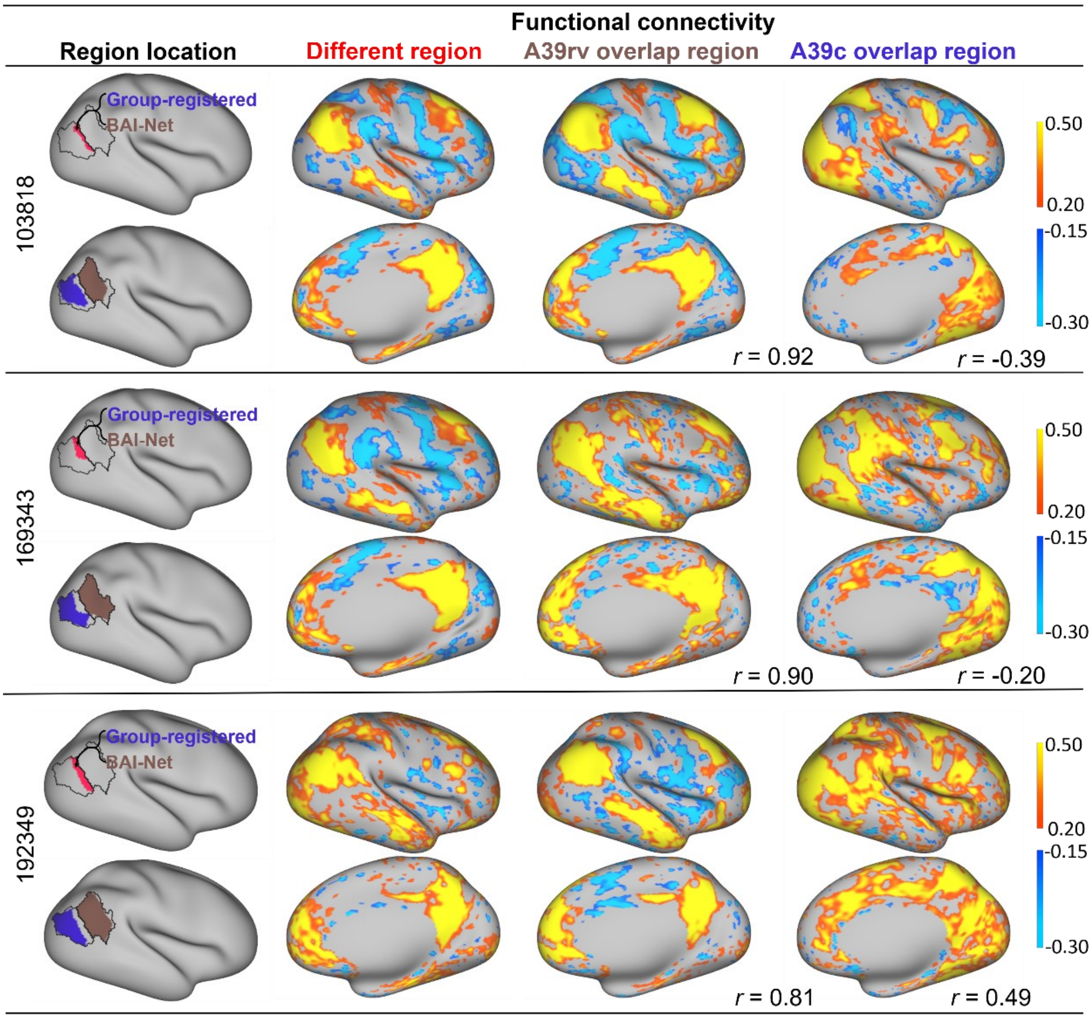


Figure S8. Functional connectivity profiles of the A39rv and A39c on two subjects from the HCP test-retest dataset. The locations of the seed regions are shown in the first column, and their functional connectivity maps are shown in the $2^{nd}$-$4^{th}$ columns. Specifically, for each subject, three seed regions were extracted based on the comparison between the registration-based atlas and the BAI-Net individualized parcellation maps, including the differential area (in red) and the common areas in A39r (in brown) and A39c (in blue). The black lines indicate the borders between A39c and A39rv in the registration-based atlas. The left three columns show the areal functional connectivity profiles for these three areas (red, brown, blue).

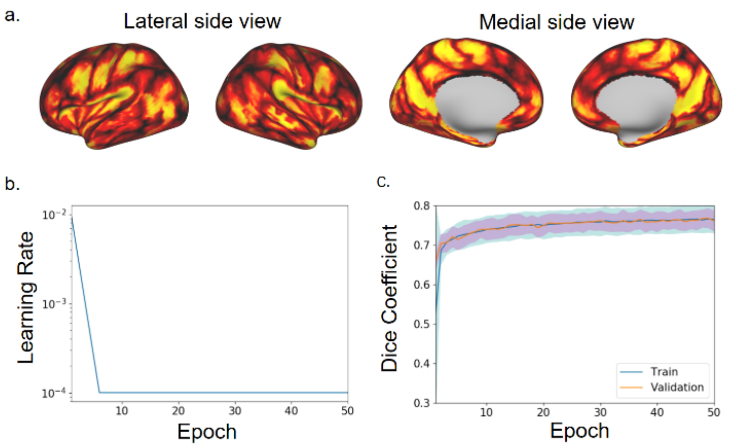


Figure S9. Training details of the graph convolution network in the left hemisphere. a: Group areal probability map used as weights in the loss function. b: learning rate with epochs. c: Overlaps between the individual atlas for the left-hemisphere model.

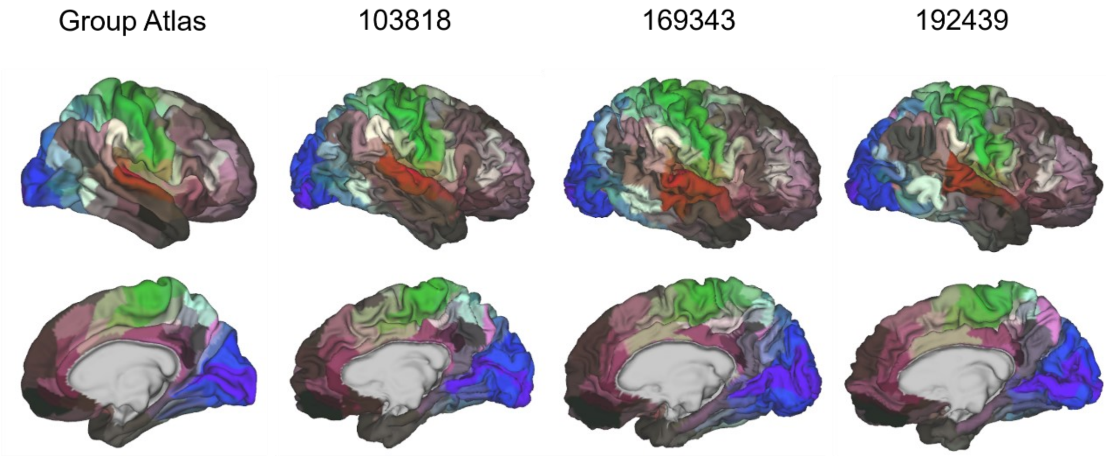


Figure S10. The implementation of the BAI-Net method on Glasser's atlas [2]. Three example subjects are shown in right hemisphere. The averaged intra-subject and inter-subject topography similarity on cartographies is Dice=0.824 and 0.673, respectively.

### Supplementary Tables

**Table S1. Names of the 72 fiber tracts**

| Index | Abbrev. | Full name | Index | Abbrev. | Full name |
|---|---|---|---|---|---|
| 0 | AF_left | Arcuate fascicle | 36 | SLF_I_right | Superior longitudinal fascicle I |
| 1 | AF_right | Arcuate fascicle | 37 | SLF_II_left | Superior longitudinal fascicle II |
| 2 | ATR_left | Anterior thalamic radiation | 38 | SLF_II_right | Superior longitudinal fascicle II |
| 3 | ATR_right | Anterior thalamic radiation | 39 | SLF_III_left | Superior longitudinal fascicle III |
| 4 | CA | Commissure anterior | 40 | SLF_III_right | Superior longitudinal fascicle III |
| 5 | CC_1 | Rostrum | 41 | STR_left | Superior thalamic radiation |
| 6 | CC_2 | Genu | 42 | STR_right | Superior thalamic radiation |
| 7 | CC_3 | Rostral body (Premotor) | 43 | UF_left | Uncinate fascicle |
| 8 | CC_4 | Anterior midbody (Primary motor) | 44 | UF_right | Uncinate fascicle |
| 9 | CC_5 | Posterior midbody (Primary somatosensory) | 45 | CC | Corpus callosum-all |
| 10 | CC_6 | Isthmus | 46 | T_PREF_left | Thalamo-prefrontal |
| 11 | CC_7 | Splenium | 47 | T_PREF_right | Thalamo-prefrontal |
| 12 | CG_left | Cingulum | 48 | T_PREM_left | Thalamo-premotor |
| 13 | CG_right | Cingulum | 49 | T_PREM_right | Thalamo-premotor |
| 14 | CST_left | Corticospinal tract | 50 | T_PREC_left | Thalamo-precentral |
| 15 | CST_right | Corticospinal tract | 51 | T_PREC_right | Thalamo-precentral |
| 16 | MLF_left | Middle longitudinal fascicle | 52 | T_POSTC_left | Thalamo-postcentral |
| 17 | MLF_right | Middle longitudinal fascicle | 53 | T_POSTC_right | Thalamo-postcentral |
| 18 | FPT_left | Fronto-pontine tract | 54 | T_PAR_left | Thalamo-parietal |
| 19 | FPT_right | Fronto-pontine tract | 55 | T_PAR_right | Thalamo-parietal |
| 20 | FX_left | Fornix | 56 | T_OCC_left | Thalamo-occipital |
| 21 | FX_right | Fornix | 57 | T_OCC_right | Thalamo-occipital |
| 22 | ICP_left | Inferior cerebellar peduncle | 58 | ST_FO_left | Striato-fronto-orbital |
| 23 | ICP_right | Inferior cerebellar peduncle | 59 | ST_FO_right | Striato-fronto-orbital |
| 24 | IFO_left | Inferior occipito-frontal fascicle | 60 | ST_PREF_left | Striato-prefrontal |
| 25 | IFO_right | Inferior occipito-frontal fascicle | 61 | ST_PREF_right | Striato-prefrontal |
| 26 | ILF_left | Inferior longitudinal fascicle | 62 | ST_PREM_left | Striato-premotor |
| 27 | ILF_right | Inferior longitudinal fascicle | 63 | ST_PREM_right | Striato-premotor |
| 28 | MCP | Middle cerebellar peduncle | 64 | ST_PREC_left | Striato-precentral |
| 29 | OR_left | Optic radiation | 65 | ST_PREC_right | Striato-precentral |
| 30 | OR_right | Optic radiation | 66 | ST_POSTC_left | Striato-postcentral |
| 31 | POPT_left | Parieto-occipital pontine | 67 | ST_POSTC_right | Striato-postcentral |
| 32 | POPT_right | Parieto-occipital pontine | 68 | ST_PAR_left | Striato-parietal |
| 33 | SCP_left | Superior cerebellar peduncle | 69 | ST_PAR_right | Striato-parietal |
| 34 | SCP_right | Superior cerebellar peduncle | 70 | ST_OCC_left | Striato-occipital |
| 35 | SLF_I_left | Superior longitudinal fascicle I | 71 | ST_OCC_right | Striato-occipital |

**Table S2. Averaged predictability of 58 individual cognitive behaviors using individual topography of IC and BAI-Net methods.**

| Description | IC | BAI-Net | P |
|---|---|---|---|
| Visual Episodic Memory | 0.158 | 0.166 | 0.142 |
| Cognitive flexibility (DCCS) | 0.115 | 0.086 | *** |
| Inhibition (Flanker task) | 0.119 | 0.092 | *** |
| Fluid Intelligence (PMAT) | 0.168 | 0.210 | *** |
| Reading (pronunciation) | 0.197 | 0.210 | 0.006 |
| Vocabulary (picture matching) | 0.196 | 0.173 | *** |
| Processing Speed | 0.013 | 0.024 | 0.047 |
| Delay Discounting | 0.095 | 0.143 | *** |
| Spatial orientation | 0.189 | 0.221 | *** |
| Sustained Attention - Sens. | -0.005 | 0.071 | *** |
| Sustained Attention - Spec. | 0.029 | 0.073 | *** |
| Verbal Episodic Memory | 0.094 | 0.117 | *** |
| Working Memory (list sorting) | 0.115 | 0.132 | *** |
| Cognitive status (MMSE) | 0.059 | 0.084 | *** |
| Sleep quality (PSQI) | -0.036 | 0.083 | *** |
| Walking endurance | 0.201 | 0.215 | *** |
| Walking Speed | -0.011 | 0.031 | *** |
| Manual dexterity | 0.175 | 0.128 | *** |
| Grip strength | 0.512 | 0.523 | *** |
| Odor identification | 0.039 | 0.087 | *** |
| Pain Interference Survey | -0.068 | -0.034 | *** |
| Taste intensity | 0.070 | 0.00 | *** |
| Contrast Sensitivity | 0.009 | -0.033 | *** |
| Emotional Face Matching | 0.079 | 0.071 | 0.091 |
| Arithmetic | 0.107 | 0.124 | 0.01 |
| Story comprehension | 0.165 | 0.180 | 0.003 |
| Relational processing | 0.224 | 0.234 | 0.035 |
| Social Cognition - random | -0.014 | 0.115 | *** |
| Social Cognition - interaction | -0.009 | 0.060 | *** |
| Working Memory (n-back) | 0.191 | 0.200 | *** |
| Agreeableness (NEO) | 0.125 | 0.162 | *** |
| Openness (NEO) | 0.065 | 0.093 | *** |
| Conscientiousness (NEO) | 0.097 | 0.032 | *** |
| Neuroticism (NEO) | 0.049 | 0.037 | 0.131 |
| Extraversion (NEO) | 0.004 | 0.110 | *** |
| Emot. Recog. - Total | 0.089 | 0.151 | *** |
| Emot. Recog. - Angry | 0.054 | 0.083 | *** |
| Emot. Recog. - Fear | 0.018 | 0.057 | *** |
| Emot. Recog. - Happy | -0.053 | 0.047 | *** |
| Emot. Recog. - Neutral | 0.020 | 0.045 | 0.002 |
| Emot. Recog. - Sad | -0.016 | 0.110 | *** |
| Anger - Affect | 0.025 | 0.077 | *** |
| Anger - Hostility | 0.036 | 0.050 | 0.018 |
| Anger - Aggression | 0.193 | 0.141 | *** |
| Fear - Affect | 0.029 | 0.058 | *** |
| Fear - Somatic Arousal | 0.118 | 0.042 | *** |
| Sadness | 0.027 | 0.084 | *** |
| Life Satisfaction | 0.075 | 0.125 | *** |
| Meaning & Purpose | 0.026 | 0.031 | 0.377 |
| Positive Affect | 0.061 | 0.102 | *** |
| Friendship | 0.055 | 0.033 | *** |
| Loneliness | -0.035 | 0.011 | *** |
| Perceived Hostility | -0.017 | 0.028 | *** |
| Perceived Rejection | 0.018 | 0.081 | *** |
| Emotional Support | 0.109 | 0.162 | *** |
| Instrument Support | 0.000 | 0.007 | 0.224 |

| | | | |
|---|---|---|---|
| Perceived Stress | 0.068 | 0.071 | 0.719 |
| Self-Efficacy | 0.139 | 0.116 | *** |

* Indicates the significant ($p$<0.001) between two methods in the two-sample t test. The gray rows are the 31 selected cognition behaviors significantly ($p$<0.001) predicted in at least one trial.
